# The Role of Osteocyte Estrogen Receptor β in Bone Mass Maintenance and Tibial Stiffness Following Sex Hormone Withdrawal in Male and Female Mice

**DOI:** 10.64898/2026.05.31.729157

**Authors:** Xiaoyu Xu, Mitchell Hoge, Janiel A. Chin-Tai, Russell P. Main

## Abstract

Sex hormones are essential regulators of skeletal maintenance, but the cell-specific mechanisms by which osteocytes mediate the skeletal consequences of sex hormone deficiency remain incompletely understood. Osteocyte estrogen receptor β (Ot-ERβ) has been implicated in sex-specific regulation of bone mass, particularly in male mice, yet its role in coordinating bone morphology and mechanical competence following sex hormone withdrawal is unclear. In this study, male and female mice with osteocyte-targeted ERβ deletion (ERβ-dOT) and littermate controls were subjected to orchiectomy (ORX), ovariectomy (OVX), or sham surgery at 20 weeks of age. Four weeks later, vertebral and tibial bone morphology were assessed by micro-computed tomography, and tibial mechanical behavior was evaluated using strain gauge-calibrated, microCT-based finite element modeling. ORX induced substantial cancellous bone loss in the lumbar vertebra and proximal tibia of male mice regardless of genotype. In cortical bone, however, ORX reduced tibial cortical area and minimum moment of inertia in male littermate controls, whereas these cortical deficits were attenuated in male ERβ-dOT mice. Consistent with these morphological changes, ORX increased finite element-predicted peak tensile and compressive strains in tibial cortical and cancellous compartments and reduced whole-bone stiffness in male controls, but these mechanical deteriorations were largely prevented by Ot-ERβ deletion. In contrast, OVX produced modest changes in female tibial cortical geometry and increased cancellous bone strains, but these responses were not strongly dependent on Ot-ERβ. Together, these findings reveal that Ot-ERβ mediates the skeletal response to sex hormone withdrawal in a sex- and compartment-dependent manner. Specifically, Ot-ERβ contributes to ORX-induced deterioration of tibial cortical morphology and mechanical competence in male mice, whereas it is largely dispensable for OVX-induced skeletal changes in female mice. This work highlights osteocyte ERβ as a sex-specific regulator linking hormonal status, bone morphology, and load-induced strain environments.

## Introduction

Sex hormones are key regulators of skeletal growth and maturation, resulting in sexual dimorphism of skeletal geometry [1–4]. Age-related bone loss in humans, especially in women during and after menopause, is recognized as a result of imbalanced bone turnover caused by the declines in the sex hormones, particularly estrogen levels [5,6]. Estrogen, mainly estradiol, protects male and female skeletons by inhibiting bone resorption [7–10] and preserving bone formation [11–15] through binding to estrogen receptors (ERs). Androgen, specifically testosterone, is important for the development and remodeling of the male skeleton by repressing bone turnover and stimulating bone formation [3,4,16]. Besides binding to the androgen receptor (AR), testosterone can interact with ERs directly through ERs/AR heterodimers or indirectly through aromatization into estrogen [15,17,18], which suggests that testosterone may regulate skeletal metabolism through ERs activation. Osteocyte-ERβ (Ot-ERβ) was shown in our previous study to mediate skeletal maintenance in male but not female mice, revealing a different role of Ot-ERβ in bone mass protection between males and females [19]. However, the interactions of sex hormones and Ot-ERβ in this sex-specific regulation of skeletal maintenance remain unclear.

Gonadectomized (ORX/OVX) rodents are commonly used for investigating the effect of sex hormone deficiency on bone turnover since they follow similar bone loss patterns as aging men and women[20–27]. ERs are shown to be required for the sex hormone-mediated skeletal maintenance in several ORX/OVX studies using global ERs knockout (KO) mice [18,25,28]. Furthermore, ERα has been indicated to mediate the estrogen, but not testosterone, effects on skeletal protection in male and female mice [25,28–30]. Particularly, it is shown in a previous OVX study that Ot-ERα primarily mediates the cancellous bone preservation by estrogen but might not play a similar role in cortical bone [31], suggesting the effect of estrogen on cortical bone protection might require Ot-ERβ. Moreover, OVX-induced cancellous bone loss was partially recovered by estrogen administration in female mice lacking Ot-ERα [31], indicating that Ot-ERβ is involved in the protective effect of estrogen on cancellous bone in female mice. However, no direct evidence elucidates the role of Ot-ERβ in sex hormone regulations of female mice skeletons, and relevant studies in male mice are lacking. In addition, sex hormone deficiency-induced bone loss with aging alters bone morphology and geometry, resulting in corresponding changes in the strain environment of cortical and cancellous bone and ultimately affecting whole-bone mechanical strength [2,4,32]. However, the relevant roles of sex hormones and Ot-ERβ in mediating skeletal strength (or strain) have not been addressed in male or female mice.

This study examined bone morphology and skeletal strength in gonadectomized (ORX or OVX) young adult male and female mice with Ot-ERβ deletion by micro-CT analysis and finite element (FE) modeling, respectively. By elucidating the effect of sex hormone withdrawal on bone morphology and skeletal strength, as well as the role of Ot-ERβ in the sex hormone regulations of the skeletons in male and female mice, we hope to offer insight into the mechanism of Ot-ERβ in bone loss and skeletal weakness induced by sex hormone declines, which could potentially benefit further exploration of nonpharmacologic therapies to prevent age-related osteoporosis and associated fractures.

## Materials and Methods

### Animal model generation

Mice with Ot-ERβ deletion (ERβ-dOT) were generated by breeding mice with a floxed ERβ gene (Esr2) to Dmp1-8kb-Cre mice [33,34]. The offspring of this cross were bred to generate male and female mice with ERβ-dOT (ERβ^Cre/+; fl/fl^) and littermate controls (ERβ^−/−; fl/fl^). Using the Dmp1-8kb-Cre model system, we generated mice with complete ERβ-dOT, whereby the expression of DMP1 promotes the Cre-induced ERβ exon 3 deletion in the skeleton. While DMP1 is also expressed in late/mature osteoblasts in the mammalian skeleton, previous studies have implicated that the mature osteoblasts that produce DMP-1 are destined to differentiate into osteocytes [35]. Primers for detecting the floxed ERβ were 5’-GCATAGCGCAGTTGGTAGAG-3’ (ERβ exon 3), 5’-CTTCTTAGAGGTACGGATCCCAGCCC-3’ (ERβ exon 3), and for the Cre transgene 5’-CATCGCTCGACCAGTTTAGTTACC-3’ (Cre), and 5’-CATACCTGGAAAATGCTTCTGTCC-3’ (Cre). Mice were housed 3 to 5 per cage with 12:12 light: dark cycle and ad libitum access to food and water. All experimental procedures were approved by Purdue University’s Animal Care and Use Committee (IACUC #1203000623).

### Orchiectomy (ORX) and ovariectomy (OVX) surgeries

Male and female mice with ERβ-dOT (KO) and their littermates (LC) were gonadectomized (ORX or OVX; O) or sham-gonadectomized (Sham; S) at 20 wks of age. There are eight groups of mice in this study: MLCS, MLCO, MKOS, MKOO, FLCS, FLCO, FKOS, and FKOO (n = 7-8/group). Four weeks following the surgery, at 24 wks old, in vivo loading and strain gauge measurements were applied, and then mice were sacrificed with tibiae and spines collected for later analyses.

### In vivo tibial loading and strain gauge measurements in ERβ-dOT and LC mice

At 24 wks of age, ERβ-dOT and LC mice with ORX/OVX or Sham were applied to tibial in vivo loading for the load-strain calibration. While the mouse was anesthetized, one leg at a time, a single strain gauge (EA-06-015LA-120, Micromeasurements) was aligned longitudinally to the tibia and glued to the medial surface of the tibial midshaft following the procedure described previously[36,37]. Each strain-gauged tibia of the mouse was placed in custom-built foot and knee holders attached to a material testing device (TestBench, TA Instruments, New Castle, DE, USA) and maintained by −1 N pre-load. Cyclic triangle waveform loads with increasing peak values (−4, −6, −8, −10, −12, and −15 N) were applied to each tibia at 4Hz with a 0.15 sec of symmetric loading/ unloading and a 0.10 s rest insertion between load cycles [36,38,39]. Load and strain were synchronically recorded at 2 Hz (LabChart 7.2, ADInstruments, Colorado Springs, CO, USA). The duration of each load trial was dependent upon the time required to reach repeatable peak strains (typically < 30 s). After the in vivo loading, mice were sacrificed by CO_2_ with tibiae and spines collected. The lead wires to the gauge were cut, and the strain gauge was preserved intact on the tibia. Four consecutive load cycles were chosen for each load level to calculate the peak load-strain relationship at the gauge site of each tibia [36]. Data from the left and right tibiae were averaged for each mouse before group means were calculated. A single load-strain relationship was generated for each mouse to measure the stiffness slope. The average stiffness at the gauge site was then calculated for each group and defined as ‘gauge-based experimental stiffness’ (**exp-stiffness**).

### Micro-CT imaging and analyses

Following the in vivo loading, the spine and gauge-attached tibiae dissected from each mouse were scanned by micro-CT with an isotropic voxel resolution of 10 μm (μCT40, Scanco Medical AG, Brüttisellen, Switzerland; 55kVp, 145 mA, 300 ms integration time, no frame averaging). Volumes of interest (VOIs) for the diaphyseal cortical bone of the tibia spanned 2.5% of the average bone length centered at the mid-diaphysis (50%) and 37% of the tibial length from the proximal end [40]. Cancellous bone VOIs included the central body of the 4^th^ lumbar vertebra (L4) and the tibial metaphysis, which was measured at 5% of the average tibial length [41]. Cortical-based and cancellous-based threshold values were determined for bone tissue segmentation using the cancellous and cortical VOIs [42,43]. Threshold values used to segment the cortical and cancellous compartments from soft tissues were 357 mg HA/cm^3^ and 224 mg HA/cm^3^, respectively, irrespective of sex, genotype, or surgery, and visualized by eye [44].

Measured parameters for cortical bone analysis (for tibia only) included cortical bone mass density (Ct. BMD; mg HA/mm^3^), cortical bone area (Ct. Ar; mm^2^), total area (T. Ar; mm^2^), bone marrow area (Mar. Ar; mm^2^), maximum and minimum moments of inertia (for tibiae only; Imax, Imin; mm^4^), and cortical bone tissue mineral density (for tibiae only, Ct. BMD, mg HA/mm^3^). Measurements for cancellous bone analysis (for both L4 body and tibia) included total tissue volume (Tb. TV; mm^3^), trabecular bone volume (Tb. BV; mm^3^), bone volume fraction (Tb. BV/TV; %), trabecular number (Tb. N; 1/mm), trabecular thickness (Tb. Th; mm), trabecular separation (Tb. Sp; mm), trabecular bone tissue mineral density (Tb. BMD, mg HA/mm^3^)[45].

Based on the outlier analysis of the body mass, mice at 24 wks of age with overweight or less weight were excluded from the group for the bone morphology analysis. Specifically, 2 MLCS, 1 FKOO, 1 FKOS, 2 FLCO, and 1 FLCS excluded.

### Micro-CT based finite element analysis (FEA)

FE analysis was performed using the micro-CT scans, stated in the above section, to characterize the full-field strain environments throughout the tibiae following methods outlined in our previous studies (Abaqus 6.23, Simulia, MI, USA) [37,41]. Briefly, the micro-CT images of each tibia were processed by a MATLAB program to generate a three-dimensional (3D) FE mesh model consisting of tetrahedral elements. Heterogeneous material properties were assigned to the FE model based on the grayscale values of micro-CT images. Three-dimensional tibia models from different groups were assigned to the same sets of loading (−7N) and boundary conditions in the FE analysis to replicate the in vivo tibial compressive loading configuration and the subsequent statistical comparison. Comprehensive sensitivity analyses were performed to determine the optimized model parameters to match the predicted strains with gauge-measured strain at the medial surface of the tibial midshaft [37].

The FE-predicted strain at the gauge site was validated by averaging the nodal strains aligned with the longitudinal axis of the overlying strain gauge. FE-predicted stiffness (FE-stiffness, N/με) was calculated by taking the applied load (−7N) over the FE-predicted strain at the gauge site (Supplementary Table S2). The FE-predicted whole-bone stiffness (N/mm) was defined as the applied compressive load over displacement along the loading axis. The strain environments for the diaphyseal cortical and metaphyseal cancellous bone of the gauged tibiae were also analyzed using FE and compared across genotype and surgery. The cortical and cancellous VOIs presented for the bone morphology analyses correspond to the VOIs used in the micro-CT analyses. Cancellous-based and cortical-based threshold values were determined for bone tissue segmentation using the cancellous and cortical VOIs [42,43]. Consistent with our previous studies, principal peak strains were defined using the cut-off values for the upper 95th percentile of the peak maximum (tensile) or minimum (compressive) principal strains for each VOI, by which to eliminate anomalous high-strain elements and numerical errors that arose during FE modeling [41,46]. Samples were excluded from the FE analysis because of (1) the broken fibulae during postmortem dissection, (2) the detached strain gauge from the bone before scanning by micro-CT, and (3) the over or less body mass compared to the other mice within the same group. The final sample size of each group for the FEA is n = 5 for each male mouse group (MLCS, MLCO, MKOS, and MKOO) and n = 4-6 for the female mice group (FLCS, FLCO, FKOS, and FKOO).

The FE model used in this study was validated with the in vivo strain gauge measurements by Student’s paired t-test (p < 0.05). The FE-predicted strain at the gauge site matched the strain measured by in vivo gauge for each genotype-surgery recombination (Supplementary Table S3). Also, the gauge-site stiffness predicted by FE and measured by in vivo gauge showed no difference, regardless of sex, genotype, and surgery (Supplementary Table S2).

### Statistical analysis

Data are presented as means ± SD. Assumptions of normality and homogeneity of variance were tested by running the Shapiro-Wilk test and Levene’s test, respectively, before statistical analyses. Differences in morphological and mechanical parameters (morphology, stiffness, or strain) between genotypes (LC vs. KO) or surgery (ORX/ORX vs. Sham) were analyzed via Student’s t-test (p < 0.05). The main effects of genotype and surgery and the genotype-surgery interaction on the morphology and skeletal strength were determined by two-way ANOVA analysis. For significant genotype-surgery interaction (p < 0.05), subsequent pairwise comparisons between genotypes and/or surgery were tested using Bonferroni correction. All results presented are significant unless otherwise stated. If no significant interaction was present, only the main effects were reported. Percentage differences were calculated for the effect of genotype or surgery as [(ERβ-dOT - LC) * 100 / LC] or as [(Surgery group – Sham group) * 100 / Sham group], respectively. Comparisons between sexes for most outcome measures were not conducted in this study. All statistical tests were run by SPSS Statistics 28 (IBM Inc., Chicago, IL, USA).

## Results

### Mouse characteristics

At 24 wks of age, body mass increased in female mice with the OVX surgery for both ERβ-dOT (+10.9%) and LC (+8.4%) relative to the corresponding Sham mice (Supplementary Table S1). However, decreased body weight appeared after the ORX surgery in male ERβ-dOT (−2.0%) mice at 24 wks of age compared to the Sham group. In contrast, male LC mice with ORX kept a similar body weight as the Sham group (Supplementary Table S1). The changes in body weight in gonadectomized male and female mice might be caused by the varied body fat mass due to the sex hormone withdrawal [47,48].

### Cortical and cancellous bone morphology in gonadal intact male and female ERβ-dOT and LC mice

Ot-ERβ was shown in previous studies to mediate cortical and cancellous bone regulation differently by sex [19]. Gonadal intact male mice appeared to have a decreased cortical bone mass with Ot-ERβ deletion, demonstrated by the reduced cortical Imin (37% only: −16.8%) and the trends of decreased tibial cortical area (Ct. Ar; 37%: −4.2%, p = 0.06; 50%: −4.0%, p = 0.08) compared to the LC mice (Table 1A). No difference in tibial cortical bone, 37% or 50%, was caused by the absence of Ot-ERβ in female ERβ-dOT mice. Moreover, neither the cancellous bone morphology nor the mineral density in the L4 body and proximal tibia was altered by the deletion of Ot-ERβ for male and female mice (Table 1B).

### Ot-ERβ mediated the effects of sex hormone withdrawal differently by sex and bone compartment

The ORX-induced cancellous bone loss occurred in the L4 body and proximal tibia in male ERβ-dOT and LC mice, as demonstrated by the decreased Tb. BV/TV and Tb. N relative to the corresponding Sham group (Table 2A, Fig.1), even though the trabecular bone densities (Tb. BMD) of L4 body and proximal tibia were not altered by ORX in ERβ-dOT and LC male mice. Different from male mice, cancellous bone mass in female LC mice, L4 or proximal tibiae, was not altered by OVX-induced sex hormone deficiency (Table 2B, Fig.1). In contrast, cancellous bone loss in L4 only was observed in female ERβ-dOT mice after OVX surgery, evidenced by the significant decrease in L4 Tb. BV/TV (−18.5%) relative to the Sham. However, for both male and female mice, the effects of ORX and OVX on cancellous bone were not affected by the deletion of Ot-ERβ (Table 2, Fig.1).

**Figure 1.**
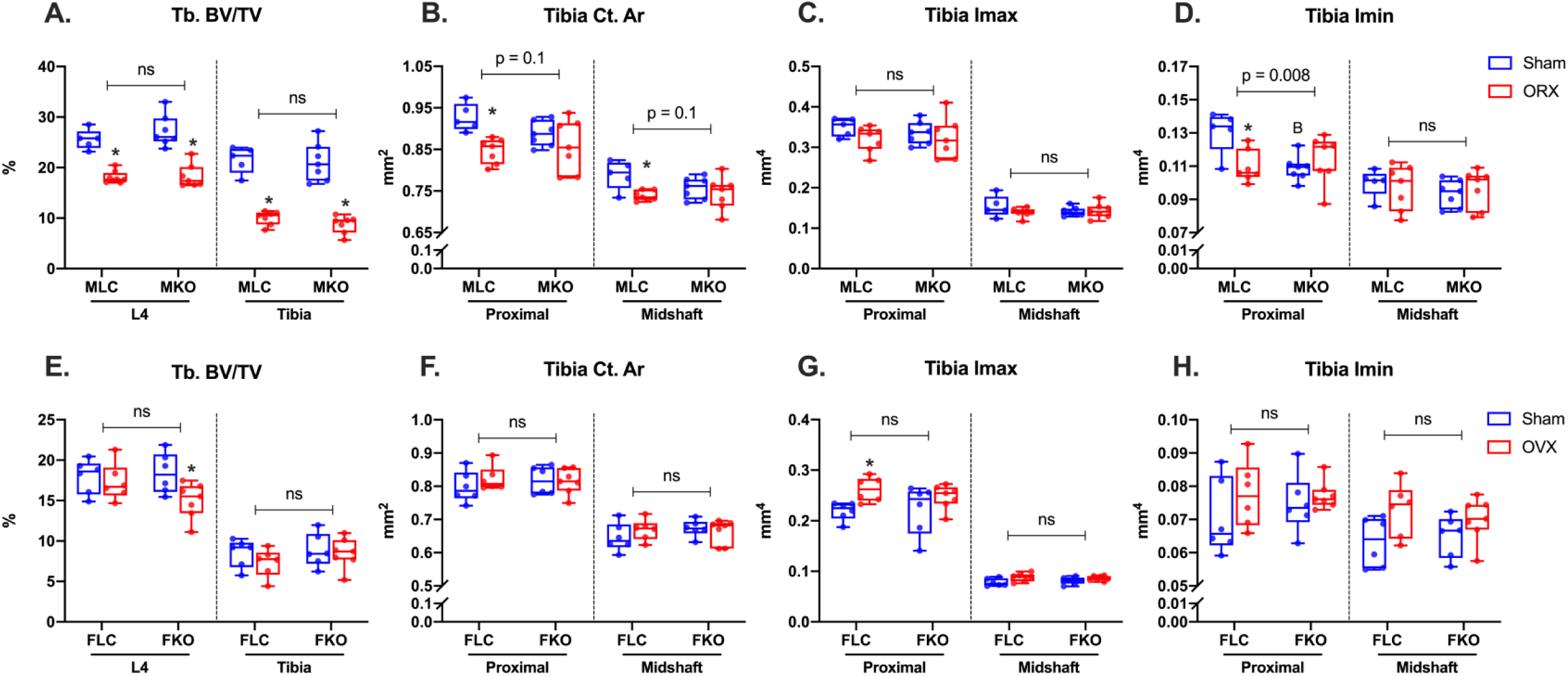
Cortical and cancellous bone morphology of vertebrae and tibiae in male and female LC and ERβ-dOT (KO) mice with gonadectomy (ORX/OVX) or Sham (Sham) surgeries. Lumbar vertebral body (L4) and tibia of male (A-D) and female (E-H) LC (MLC/FLC, blue) and ERβ-dOT (MKO/FKO, red) mice after the gonadectomy (ORX/OVX) or Sham (Sham) surgeries were analyzed by micro-CT at 24 wks of age. Trabecular bone volume fraction (Tb. BV/TV) of L4 and proximal tibia (A, E) as well as cortical area (Ct. Ar) and the maximum and minimum moments of inertia (Imax, Imin) at proximal (37%) and midshaft (50%) tibia are shown as boxplots with data points (n = 6). Asterisks ‘✽’ show the significant surgery effect (ORX/OVX versus Sham) by Student t-test (p < 0.05). ns: p > 0.1. The effects of surgery and genotype and their interaction were tested by two-way ANOVA followed by pair-wise comparisons with Bonferroni correction. Specific p-value shown when 0.001 < p < 0.1. ns: p > 0.1. Capital letter ‘B’ indicates the significant difference by genotype (ERβ-dOT versus LC) determined by two-way ANOVA with Bonferroni pair-wise comparison.

**Table 2.** Cancellous bone morphology of vertebral body (L4) and proximal tibia of male (A) and female (B) ERβ-dOT and LC mice with gonadectomy (ORX/OVX) or Sham at 24 wks of age.

| **A.**<br>**Cancellous Bone** | **MLC** |  |  | **MKO** |  |  | **2-way ANOVA Results** |  |  |
| --- | --- | --- | --- | --- | --- | --- | --- | --- | --- |
|  | Sham | ORX | Diff (%) | Sham | ORX | Diff (%) | Surgery | Genotype | Interaction |
| **L4 Body** | n = 5 | n = 8 |  | n = 7 | n = 7 |  |  |  |  |
| Tb. TV, mm<sup>3</sup> | 1.85±0.15 | 1.71±0.22 | -8.0 | 1.82±0.14 | 1.86±0.14 | 2.2 | NS | NS | NS |
| Tb. BV, mm<sup>3</sup> | 0.48±0.07 | 0.31±0.04* | -35.4 | 0.50±0.08 | 0.34±0.05* | -31.3 | 0 | NS | NS |
| Tb. BV/TV, % | 25.6±2.0 | 18.1±0.0* | -29.4 | 27.3±3.2 | 18.4±2.3* | -32.7 | 0 | NS | NS |
| Tb. N, 1/mm | 4.4±0.2 | 3.6±0.2* | -18.7 | 4.5±0.5 | 3.6±0.3* | -19.5 | 0 | NS | NS |
| Tb. Th, mm | 0.057±0.001 | 0.056±0.002 | -2.9 | 0.059±0.002 | 0.056±0.003* | -5.4 | 0.01 | NS | NS |
| Tb. Sp, mm | 0.21±0.01 | 0.28±0.02* | 31.0 | 0.22±0.03 | 0.28±0.02* | 24.4 | 0 | NS | NS |
| Tb. BMD, mg HA/mm<sup>3</sup> | 772.6±22.5 | 755.2±15.3 | -2.3 | 759.9±24.9 | 756.2±19.7 | -0.5 | NS | NS | NS |
| **Proximal Tibia** |  |  |  |  |  |  |  |  |  |
| Tb. TV, mm<sup>3</sup> | 2.46±0.12 | 2.34±0.17 | -4.8 | 2.22±0.16 | 2.36±0.07* | 6.7 | NS | NS | NS |
| Tb. BV, mm<sup>3</sup> | 0.53±0.08 | 0.24±0.04* | -55.6 | 0.47±0.10 | 0.21±0.04* | -56.2 | 0 | NS | NS |
| Tb. BV/TV, % | 21.6±2.6 | 10.0±1.4* | -53.4 | 21.1±3.8 | 8.7±1.7* | -58.7 | 0 | NS | NS |
| Tb. N, 1/mm | 4.8±0.3 | 3.3±0.3* | -31.7 | 4.6±0.5 | 3.3±0.2* | -28.6 | 0 | NS | NS |
| Tb. Th, mm | 0.063±0.003 | 0.063±0.004 | 1.3 | 0.062±0.003 | 0.062±0.007 | 0.5 | NS | NS | NS |
| Tb. Sp, mm | 0.19±0.02 | 0.30±0.03* | 58.1 | 0.20±0.03 | 0.30±0.02* | 51.8 | 0 | NS | NS |
| Tb. BMD, mg HA/mm<sup>3</sup> | 745.5±13.5 | 760.5±19.9 | 2.0 | 751.3±22.9 | 761.8±15.5 | 1.4 | 0.10 | NS | NS |

**Table 2 Continued**
| **B.**<br>**Cancellous Bone** | **FLC** |  |  | **FKO** |  |  | **2-way ANOVA Results** |  |  |
| --- | --- | --- | --- | --- | --- | --- | --- | --- | --- |
|  | Sham | OVX | Diff (%) | Sham | OVX | Diff (%) | Surgery | Genotype | Interaction |
| **L4 Body** | n = 6 | n = 6 |  | n = 6 | n = 7 |  |  |  |  |
| Tb. TV, mm<sup>3</sup> | 1.64±0.19 | 1.90±0.12 | 15.9 | 1.48±0.14 | 1.79±0.13* | 21.2 | 0 | NS | NS |
| Tb. BV, mm<sup>3</sup> | 0.29±0.03 | 0.33±0.05 | 12.5 | 0.27±0.04 | 0.27±0.05 | -0.5 | NS | NS | NS |
| Tb. BV/TV, % | 18.0±2.1 | 17.3±2.3 | -3.9 | 18.4±2.5 | 15.0±2.2* | -18.5 | 0.004 | NS | NS |
| Tb. N, 1/mm | 3.0±0.3 | 2.9±0.3 | 0.5 | 2.9±0.2 | 2.8±0.2* | -7.8 | 0.07 | NS | NS |
| Tb. Th, mm | 0.060±0.002 | 0.056±0.002* | -6.8 | 0.061±0.002 | 0.054±0.003* | -12.1 | 0 | NS | NS |
| Tb. Sp, mm | 0.34±0.04 | 0.34±0.04 | -1.0 | 0.34±0.02 | 0.37±0.04* | 9.2 | 0.07 | NS | NS |
| Tb. BMD, mg HA/mm<sup>3</sup> | 773.3±21.6 | 762.5±18.3 | -1.4 | 776.0±30.2 | 753.9±25.9 | -2.8 | 0.06 | NS | NS |
| **Proximal Tibia** |  |  |  |  |  |  |  |  |  |
| Tb. TV, mm<sup>3</sup> | 1.79±0.13 | 2.02±0.20* | 13.2 | 1.74±0.10 | 2.01±0.12* | 16.0 | 0 | NS | NS |
| Tb. BV, mm<sup>3</sup> | 0.15±0.03 | 0.15±0.05 | -0.9 | 0.15±0.04 | 0.17±0.04 | 10.6 | NS | NS | NS |
| Tb. BV/TV, % | 8.5±1.7 | 7.3±1.7 | -14.2 | 8.9±2.1 | 8.5±1.9 | -3.9 | NS | NS | NS |
| Tb. N, 1/mm | 2.7±0.3 | 2.5±0.3 | -8.8 | 3.0±0.4 | 2.5±0.2* | -14.6 | 0.01 | NS | NS |
| Tb. Th, mm | 0.070±0.004 | 0.066±0.009 | -5.8 | 0.063±0.006<sup>A</sup> | 0.069±0.007 | 8.3 | NS | NS | NS |
| Tb. Sp, mm | 0.37±0.05 | 0.41±0.05 | 10.7 | 0.34±0.04 | 0.39±0.03* | 17.0 | 0.01 | NS | NS |
| Tb. BMD, mg HA/mm<sup>3</sup> | 811.5±15.6 | 784.2±25.7 | -3.4 | 785.7±25.2 | 786.7±25.2 | 0.1 | NS | NS | NS |

Note: Data are presented as Means ± SD values (n = 5~8). Abbreviation: MLC, male LC mice; MKO, male ERβ-dOT mice; FLC, female LC mice; FKO, female ERβ-dOT mice; Tb. TV, trabecular tissue volume; Tb. BV, trabecular bone volume; Tb. BV/TV, trabecular bone ratio; Tb. N, trabecular number; Tb. Th, trabecular thickness; Tb. Sp, trabecular separation; Tb. BMD, trabecular bone mineral density. Asterisks ‘\*’ indicates the significant surgery effect (ORX/OVX versus Sham) by Student t-test (p < 0.05). Capital letter ‘A’ shows the significant difference between genotypes (ERβ-dOT versus LC) for the Sham mice by Student t-test (p < 0.05).
The effects of surgery and genotype and their interaction were tested by two-way ANOVA followed by pair-wise comparisons with Bonferroni correction. Specific p-value shown when 0.001 < p < 0.1. NS: p > 0.1. Diff (%): Difference between ORX/OVX and Sham in percentage calculated as 100\*(ORX/OVX – Sham)/Sham.

Different from the cancellous bone, a sex-dependent regulation of Ot-ERβ was observed in tibial cortical bone. When subjected to OVX, female LC mice showed a decreased cortical bone mineral density (Ct. BMD, −1.6%) at the proximal but not midshaft tibia, followed by enlarged bone marrow area (Mar. Ar, +14.1%) and increased Imax (+19.5%), compared to the Sham LC mice (Table 3B, Fig.1). In contrast, OVX did not affect the cortical bone mass in female mice lacking Ot-ERβ, although the total area (T. Ar) and bone marrow area (Mar. Ar) at proximal and midshaft tibiae were increased by OVX (Table 3B). Moreover, OVX-induced changes in cortical bone were not mediated by Ot-ERβ deletion in female mice (Table 3B).

**Table 3.** Cortical bone morphology of proximal (37%) and midshaft (50%) tibiae of male (A) and female (B) ERβ-dOT and LC mice with gonadectomy (ORX/OVX) or Sham at 24 wks of age.

| A.<br>Cortical Bone | MLC |  |  | MKO |  |  | 2-way ANOVA Results |  |  |
| --- | --- | --- | --- | --- | --- | --- | --- | --- | --- |
|  | Sham | ORX | Diff (%) | Sham | ORX | Diff (%) | Surgery | Genotype | Interaction |
| **Proximal Tibia (37%)** | n = 5 | n = 7 |  | n = 7 | n = 7 |  |  |  |  |
| Ct. BMD, mg HA/mm<sup>3</sup> | 1069.8±19.7 | 1060.3±16.5 | -0.9 | 1066.8±13.7 | 1063.2±12.1 | -0.3 | NS | NS | NS |
| Ct. Ar, mm<sup>2</sup> | 0.93±0.03 | 0.85±0.03* | -8.6 | 0.89±0.03 | 0.86±0.06 | -3.2 | 0.004 | NS | 0.1 |
| T. Ar mm<sup>2</sup> | 1.64±0.08 | 1.58±0.08 | -3.4 | 1.55±0.08 | 1.58±0.10 | 2.2 | NS | NS | NS |
| Mar. Ar mm<sup>2</sup> | 0.71±0.07 | 0.74±0.09 | 3.4 | 0.66±0.06 | 0.72±0.09 | 9.4 | NS | NS | NS |
| Imin, mm<sup>4</sup> | 0.13±0.01 | 0.11±0.01* | -16.1 | 0.10±0.01<sup>B</sup> | 0.11±0.02 | 5.0 | 0.1 | 0.07 | 0.01 |
| Imax, mm<sup>4</sup> | 0.35±0.02 | 0.32±0.03 | -8.4 | 0.34±0.03 | 0.32±0.05 | -3.6 | NS | NS | NS |
| **Midshaft Tibia (50%)** |  |  |  |  |  |  |  |  |  |
| Ct. BMD, mg HA/mm<sup>3</sup> | 1151.8±35.6 | 1145.0±10.5 | -0.6 | 1148.6±13.9 | 1142.5±24.2 | -0.5 | NS | NS | NS |
| Ct. Ar, mm<sup>2</sup> | 0.79±0.04 | 0.74±0.01* | -6.3 | 0.76±0.03 | 0.74±0.04 | -1.6 | 0.01 | NS | 0.1 |
| T. Ar mm<sup>2</sup> | 1.34±0.08 | 1.33±0.09 | -0.7 | 1.30±0.05 | 1.34±0.08 | 2.8 | NS | NS | NS |
| Mar. Ar mm<sup>2</sup> | 0.55±0.05 | 0.59±0.09 | 7.3 | 0.55±0.03 | 0.59±0.05 | 8.9 | NS | NS | NS |
| Imin, mm<sup>4</sup> | 0.10±0.01 | 0.10±0.01 | -2.7 | 0.09±0.01 | 0.10±0.01 | 3.3 | NS | NS | NS |
| Imax, mm<sup>4</sup> | 0.15±0.03 | 0.14±0.01 | -9.9 | 0.14±0.01 | 0.14±0.02 | 1.7 | NS | NS | NS |

**Table 3 Continued**
| **B.**<br>**Cortical Bone** | **FLC** |  |  | **FKO** |  |  | **2-way ANOVA Results** |  |  |
| --- | --- | --- | --- | --- | --- | --- | --- | --- | --- |
|  | Sham | OVX | Diff (%) | Sham | OVX | Diff (%) | Surgery | Genotype | Interaction |
| **Proximal Tibia (37%)** | n = 6 | n = 6 |  | n = 6 | n = 7 |  |  |  |  |
| Ct. BMD, mg HA/mm<sup>3</sup> | 1111.8±9.6 | 1094.5±12.2* | -1.6 | 1118.8±17.0 | 1104.2±17.0 | -1.3 | 0.01 | NS | NS |
| Ct. Ar, mm<sup>2</sup> | 0.80±0.05 | 0.82±0.04 | 3.2 | 0.82±0.04 | 0.81±0.04 | -0.3 | NS | NS | NS |
| T. Ar mm<sup>2</sup> | 1.24±0.08 | 1.33±0.07 | 7.1 | 1.25±0.04 | 1.30±0.04* | 4.5 | 0.01 | NS | NS |
| Mar. Ar mm<sup>2</sup> | 0.44±0.04 | 0.50±0.04* | 14.1 | 0.43±0.01 | 0.49±0.05* | 13.6 | 0.001 | NS | NS |
| Imin, mm<sup>4</sup> | 0.07±0.01 | 0.08±0.01 | 10 | 0.07±0.01 | 0.08±0.00 | 3.2 | NS | NS | NS |
| Imax, mm<sup>4</sup> | 0.22±0.02 | 0.26±0.02* | 19.5 | 0.22±0.05 | 0.25±0.02 | 11.2 | 0.01 | NS | NS |
| **Midshaft Tibia (50%)** |  |  |  |  |  |  |  |  |  |
| Ct. BMD, mg HA/mm<sup>3</sup> | 1171.8±11.9 | 1163.3±8.8 | -0.7 | 1179.1±12.8 | 1164.5±19.6 | -1.2 | NS | NS | NS |
| Ct. Ar, mm<sup>2</sup> | 0.65±0.04 | 0.67±0.03 | 3.5 | 0.67±0.03 | 0.66±0.04 | -1.4 | NS | NS | NS |
| T. Ar mm<sup>2</sup> | 1.00±0.05 | 1.07±0.05* | 7.3 | 1.00±0.04 | 1.05±0.04* | 4.9 | 0.003 | NS | NS |
| Mar. Ar mm<sup>2</sup> | 0.35±0.03 | 0.40±0.03* | 14.3 | 0.33±0.04 | 0.39±0.04* | 17.9 | 0.001 | NS | NS |
| Imin, mm<sup>4</sup> | 0.06±0.01 | 0.07±0.01 | 15.2 | 0.07±0.01 | 0.07±0.01 | 7.7 | 0.02 | NS | NS |
| Imax, mm<sup>4</sup> | 0.08±0.01 | 0.09±0.01 | 12.5 | 0.08±0.01 | 0.09±0.00 | 5.7 | 0.02 | NS | NS |

Note: Data are presented as Means ± SD values (n = 5~8). Abbreviation: MLC, male LC mice; MKO, male ERβ-dOT mice; FLC, female LC mice; FKO, female ERβ-dOT mice; Ct. BMD, cortical bone mineral density; Ct. Ar, cortical bone area; T. Ar, total area; Mar. Ar, bone marrow area; Imin, minimum moments of inertia; Imax, maximum moments of inertia. Diff (%): Difference between ORX/OVX and Sham in percentage calculated as 100\*(ORX/OVX – Sham)/Sham. Asterisks ‘\*’ indicates significant surgery effect (ORX/OVX versus Sham) by Student t-test (p < 0.05). The effects of surgery and genotype and their interaction were tested by two-way ANOVA followed by pair-wise comparisons with Bonferroni correction. Specific p-value shown when 0.001 < p < 0.1. NS: p > 0.1. Capital letter ‘B’ shows the significant difference by genotype (ERβ-dOT versus LC) determined by two-way ANOVA with Bonferroni pair-wise comparison.

Differently, Ot-ERβ mediates the sex hormone deficiency-induced cortical bone loss in male mice. ORX in male LC mice led to significant decreases in tibial Ct. Ar (37%: −8.6%; 50%: −6.3%) and Imin (37% only: −16.1%) compared to the Sham mice, even though the tibial Ct. BMD (37% and 50%) was not affected by the ORX. However, with Ot-ERβ deletion, ORX did not cause any bone loss in tibial cortical bone (37% and 50%) as it did in the male LC mice (Table 3A, Fig.1). Thus, Ot-ERβ mediates sex hormone withdrawal-induced cortical bone loss in male but not female mice.

### FE-predicted peak tensile and compressive strains of tibial cortical and cancellous bone in gonadal intact male and female ERβ-dOT and LC mice

Load-induced principal peak strains in tibiae were measured by FE modeling to estimate the mechanical strengths of cortical and cancellous bone in male and female mice. Consistent with the bone morphology results, Ot-ERβ deletion affected the tibial cortical and cancellous strength differently by sex. For gonadal intact male mice, peak tensile and compressive strains in neither cancellous bone nor the cortical bone (37% or 50%) differed between genotypes (Table 4, Fig.2). In contrast, female ERβ-dOT mice showed lower peak tensile (37%: −16%; 50%: −23%) and compressive (37%: −22%; 50%: −19%) strains in tibial cortical bone than LC mice (Table 4, Fig.2), whereas similar peak strains (tensile and compressive) showed in the cancellous bone between genotypes for female mice (Table 4, Fig.3).

**Figure 2.**
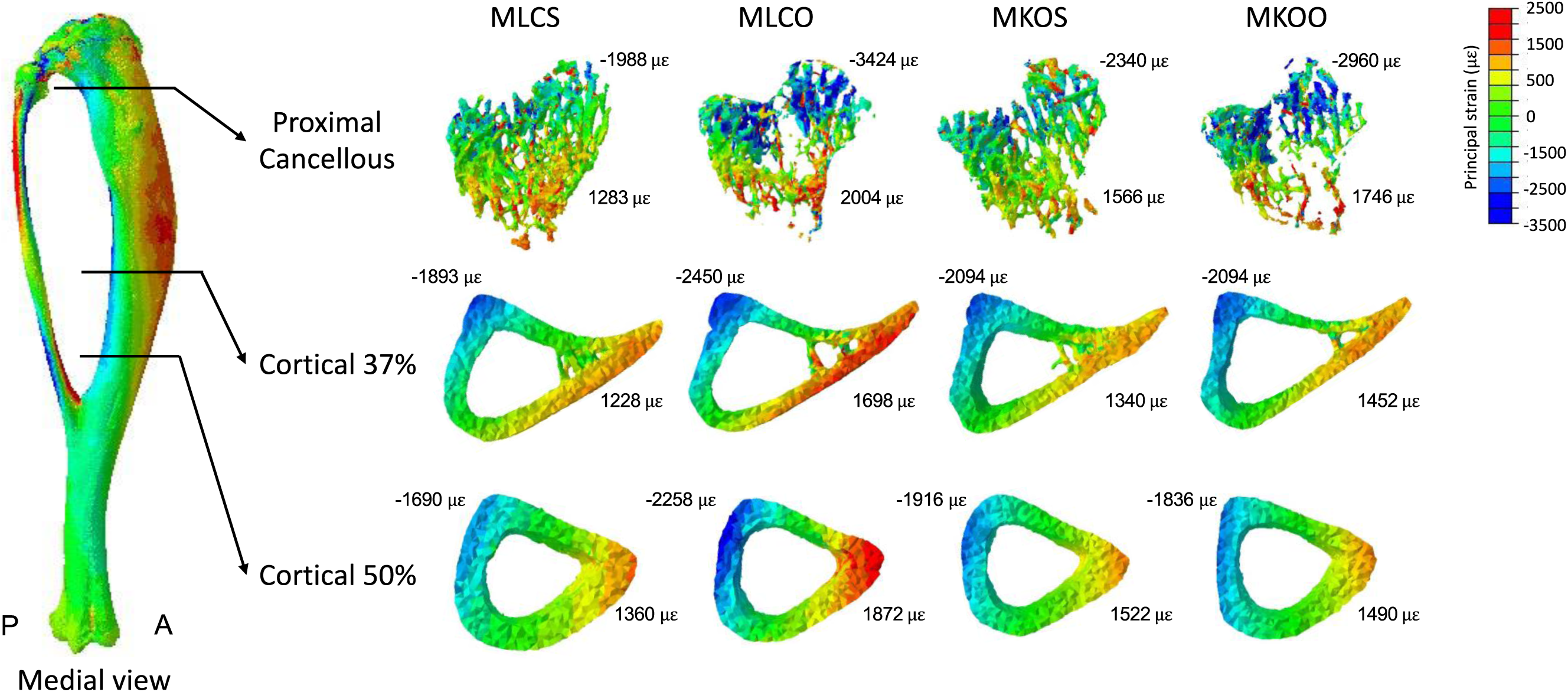
Principal strain distributions in tibial cortical and cancellous VOIs of male LC (MLC) and ERβ-dOT (MKO) mice with Sham (S) or ORX (O) surgeries. Tibiae of male LC (MLC) and ERβ-dOT (MKO) mice with Sham (MLCS/MKOS) or ORX (MLCO/MKOO) were subjected to FE-simulated compressive load of a −7N magnitude. Red and blue indicate tension and compression, respectively. Averages of 95^th^ peak maximum (tensile) and minimum (compressive) principal strains are shown by each cross-sectional image.

**Figure 3.**
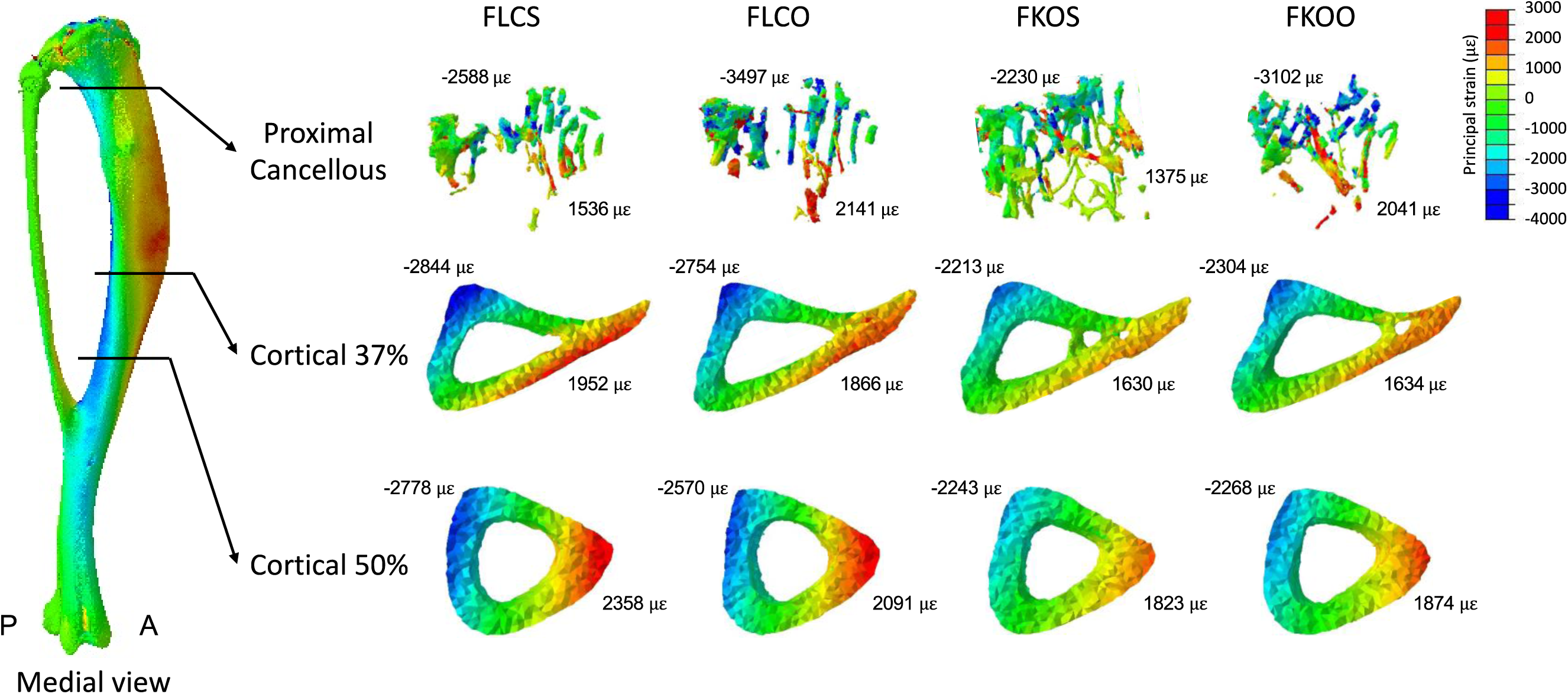
Principal strain distributions in tibial cortical and cancellous VOIs of female LC (FLC) and ERβ-dOT (FKO) mice with Sham (S) or OVX (O) surgeries. Tibiae of female LC (FLC) and ERβ-dOT (FKO) mice with Sham (FLCS/FKOS) or OVX (FLCO/FKOO) were subjected to FE-simulated compressive load of a −7N magnitude. Red and blue indicate tension and compression, respectively. Averages of 95^th^ peak maximum (tensile) and minimum (compressive) principal strains are shown by each cross-sectional image.

**Table 4.** FE-predicted peak principal strains (με) in the tibial cortical and cancellous VOIs under −7N applied load in male and female mice ERβ-dOT and LC mice with gonadectomy (ORX/OVX) or Sham.

| 24 wk<br>Male | MLC (-7N) |  | MKO (-7N) |  | 2-way ANOVA Results |  |  |
| --- | --- | --- | --- | --- | --- | --- | --- |
|  | Sham | ORX | Sham | ORX | Surgery | Genotype | Interaction |
| **CANCELLOUS** | n = 4 | n = 5 | n = 5 | n = 5 |  |  |  |
| Peak Tens | 1283±156 | 2004±142* | 1566±153<sup>B</sup> | 1746±202<sup>B</sup> | 0 | NS | 0.003 |
| Peak Comp | -1988±306 | -3424±242* | -2340±355<sup>B</sup> | -2960±608<sup>b</sup> | 0 | NS | 0.05 |
| **CORTICAL 37%** |  |  |  |  |  |  |  |
| Peak Tens | 1229±240 | 1698±110* | 1340±154 | 1452±143<sup>B</sup> | 0.002 | NS | 0.03 |
| Peak Comp | -1893±234 | -2450±218* | -2094±204 | -2094±233<sup>B</sup> | 0.02 | NS | 0.02 |
| **CORTICAL 50%** |  |  |  |  |  |  |  |
| Peak Tens | 1360±278 | 1872±127* | 1522±178 | 1490±82<sup>B</sup> | 0.01 | NS | 0.004 |
| Peak Comp | -1690±263 | -2258±157* | -1916±184 | -1836±162<sup>B</sup> | 0.01 | NS | 0.002 |
| 24 wk<br>Female | FLC (-7N) |  | FKO (-7N) |  | 2-way ANOVA Results |  |  |
|  | Sham | OVX | Sham | OVX | Surgery | Genotype | Interaction |
| **CANCELLOUS** | n = 5 | n = 5 | n = 5 | n = 5 |  |  |  |
| Peak Tens | 1536±181 | 2141±410* | 1375±118 | 2041±543* | 0.002 | NS | NS |
| Peak Comp | -2588±381 | -3497±771* | -2230±191 | -3102±788* | 0.01 | NS | NS |
| **CORTICAL 37%** |  |  |  |  |  |  |  |
| Peak Tens | 1952±303 | 1866±212 | 1630±133<sup>A</sup> | 1634±412 | NS | 0.03 | NS |
| Peak Comp | -2844±492 | -2754±398 | -2213±134<sup>A</sup> | -2304±412 | NS | 0.01 | NS |
| **CORTICAL 50%** |  |  |  |  |  |  |  |
| Peak Tens | 2358±296 | 2091±184 | 1823±168<sup>A</sup> | 1874±485 | NS | 0.01 | NS |
| Peak Comp | -2778±327 | -2570±313 | -2243±129<sup>A</sup> | -2268±454 | NS | 0.01 | NS |

Note: Data shown as Means ± SD (n = 4–5 for male mice; n = 5 for female mice). Peak Ten, the upper 95<sup>th</sup> percentile of the peak maximum (tensile) principal strains; Peak Comp, the upper 95<sup>th</sup> percentile of the peak minimum (compressive) principal strains. Asterisks ‘\*’ show the significant surgery effect by Student *t*-test ( $p < 0.05$ ). Capital letter ‘A’ indicates the significant difference between genotypes (ERβ-dOT versus LC) for the Sham mice by Student *t*-test ( $p < 0.05$ ). The effects of surgery and genotype and their interaction were tested by two-way ANOVA ( $p < 0.05$ ) followed by pair-wise comparisons with Bonferroni correction. Specific *p*-value shown when $0.001 < p < 0.1$ . NS: $p > 0.1$ . Capital letter ‘B’ shows the significant difference ( $p < 0.05$ ) by genotype by two-way ANOVA with Bonferroni pair-wise comparison. Lowercase ‘b’, $0.05 < p < 0.1$ , shows the trend of genotype effect by two-way ANOVA with Bonferroni comparison.

### Gonadectomy-induced changes in the load-induced peak strains in tibial cortical and cancellous bone were differently regulated by Ot-ERβ in male and female mice

Sex hormone withdrawal by gonadectomy surgeries caused an opposite effect on tibial bone strength between male and female mice. ORX in male LC mice led to a great decrease in tibial strength, demonstrated by the increased peak strains (tensile and compressive) in tibial cancellous and cortical bone relative to the Sham LC mice (Table 4, Fig.2). However, such ORX-induced increases in tibial strains were prevented by the deletion of Ot-ERβ in male mice, evidenced by the similar cortical and cancellous bone strains between ORX ERβ-dOT and Sham mice (Table 4, Fig.2).

In female mice, OVX-induced increases in peak tensile and compressive strains in the tibial cancellous bone were similar between ERβ-dOT and LC mice. Cortical strains (tensile and compressive) in both female ERβ-dOT and LC mice were not affected by OVX (Table 4, Fig.3). Ot-ERβ regulates the tibial whole-bone stiffness oppositely and is associated with the sex hormones differently in male and female mice

By running the FE modeling, the tibial whole-bone stiffness was estimated in male and female mice by taking the applied load (−7 N) over the whole-bone displacement. For the Sham mice, tibial whole-bone stiffness was altered differently by the deletion of Ot-ERβ in male and female mice, that it decreased in the male ERβ-dOT mice but was not altered in the female ERβ-dOT mice compared to their corresponding LC mice (Table 5).

**Table 5.**
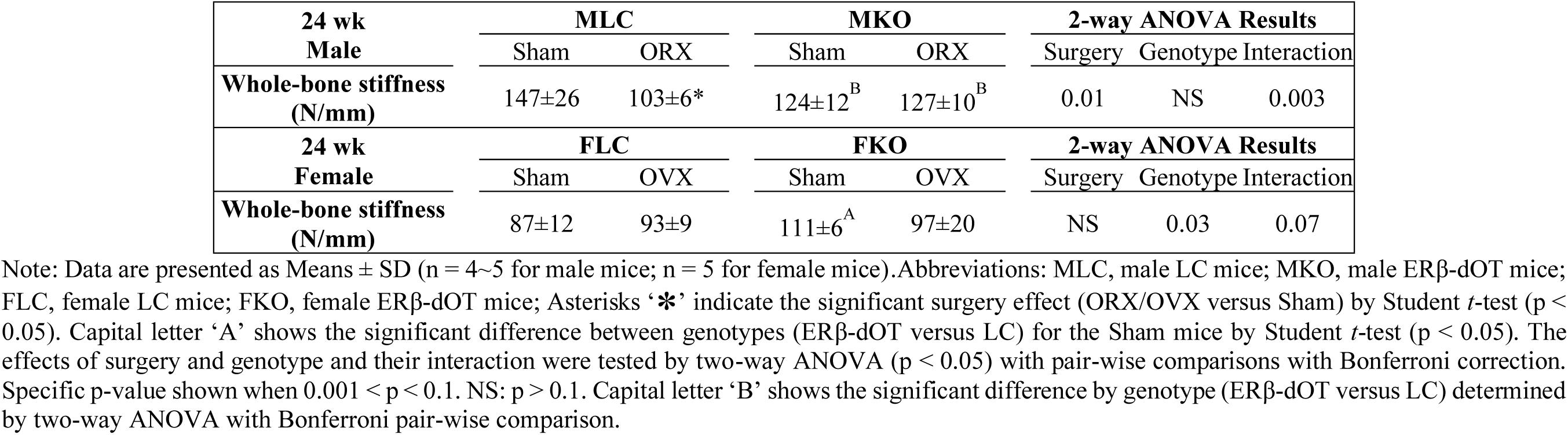
FE-predicted whole-bone stiffness under −7N applied load in male and female mice ERβ-dOT and LC mice with gonadectomy (ORX/OVX) or Sham.

| 24 wk<br>Male | MLC |  | MKO |  | 2-way ANOVA Results |  |  |
| --- | --- | --- | --- | --- | --- | --- | --- |
|  | Sham | ORX | Sham | ORX | Surgery | Genotype | Interaction |
| Whole-bone stiffness<br>(N/mm) | 147±26 | 103±6* | 124±12<sup>B</sup> | 127±10<sup>B</sup> | 0.01 | NS | 0.003 |
| 24 wk<br>Female | FLC |  | FKO |  | 2-way ANOVA Results |  |  |
|  | Sham | OVX | Sham | OVX | Surgery | Genotype | Interaction |
| Whole-bone stiffness<br>(N/mm) | 87±12 | 93±9 | 111±6<sup>A</sup> | 97±20 | NS | 0.03 | 0.07 |

Note: Data are presented as Means ± SD (n = 4~5 for male mice; n = 5 for female mice). Abbreviations: MLC, male LC mice; MKO, male ERβ-dOT mice; FLC, female LC mice; FKO, female ERβ-dOT mice; Asterisks ‘\*’ indicate the significant surgery effect (ORX/OVX versus Sham) by Student *t*-test (p < 0.05). Capital letter ‘A’ shows the significant difference between genotypes (ERβ-dOT versus LC) for the Sham mice by Student *t*-test (p < 0.05). The effects of surgery and genotype and their interaction were tested by two-way ANOVA (p < 0.05) with pair-wise comparisons with Bonferroni correction. Specific p-value shown when 0.001 < p < 0.1. NS: p > 0.1. Capital letter ‘B’ shows the significant difference by genotype (ERβ-dOT versus LC) determined by two-way ANOVA with Bonferroni pair-wise comparison.

When subjected to ORX, the tibia stiffness of male LC mice was significantly decreased relative to the Sham mice, which was consistent with the decreased cortical bone morphology (37%, Table 3) and the increased load-induced peak strains in both tibial cancellous and cortical bone (Table 4) relative to the Sham LC mice. However, Ot-ERβ deletion in male mice mitigates tibia stiffness reduction induced by sex hormone deficiency (Table 5). Unlike male mice, tibia stiffness was not altered by OVX in female LC mice, but a trend of decreased tibia stiffness (interaction, p = .07) was induced by OVX in female ERβ-dOT mice (Table 5).

## Discussion

In this study, we investigated the role of Ot-ERβ in sex hormone withdrawal-induced changes in bone morphology and skeletal strength in young adult mice. Male and female ERβ-dOT and LC mice were subjected to gonadectomy surgeries (ORX or OVX) at 20 wks of age. Four weeks after the surgery, at 24 wks old, in vivo compressive loading was applied to gauge-attached tibiae for load-strain calibration. Bone morphology was analyzed in the L4 vertebral body and tibia by micro-CT. The load-induced strains in tibial cortical and cancellous bone and the tibial whole-bone strength were simulated by FE modeling for male and female mice at 24 wks of age.

Our previous study on adult mice (30wk) suggested that Ot-ERβ inhibits vertebral bone loss in adult male mice by suppressing osteoclast-related bone resorption, but plays a minor role in adult female mice skeletons. This sexually dimorphic regulation of Ot-ERβ was also observed in young adult mice (24wk). Decreased tibial cortical Imin (37%) and Ct. Ar (37% and 50%) appeared in gonadal intact male mice with Ot-ERβ deletion compared to the LC mice, whereas no difference in cortical bone (37% and 50%) was observed in female mice between genotypes. Thus, Ot-ERβ plays roles in maintaining cortical bone morphology in young adult male but not female mice. In addition, Ot-ERβ may play a dispensable role in cancellous bone maintenance for both male and female mice, which was reflected by the similar bone morphology and density (L4 and tibia) between ERβ-dOT and LC mice. Collectively, Ot-ERβ mediates bone morphology differently in young adult mice by bone compartment and sex, such that it is required for cortical but not cancellous bone protection for male mice but may not mediate cortical and cancellous bone maintenance for female mice.

It is known that estrogens, as well as testosterone, play important roles in skeletal protection for both humans and rodents [1,15,18,49,50]. Age-related decline in circulating sex hormones is one of the primary causes of the rapid bone loss in males and females, particularly postmenopausal women [1,2,51,52]. In our study, the loss of bone mass occurred in gonadectomized male and female mice due to sex hormone deficiency. Compared to the gonadal intact (Sham) mice, ORX-induced bone mass loss appeared in male LC mice, reflected in the decreased bone morphology in both cancellous bone (L4 and tibia) and cortical bone (tibia, Table 3A). Similarly, female LC mice developed osteopenia in cortical bone after OVX, manifested by decreased bone mineral density and increased Imax at the proximal tibia. Cancellous bone (L4 and tibia) in female LC mice showed a slight decrease in OVX with no significant difference compared to the Sham. According to the ORX/OVX-induced bone loss in LC mice, sex hormones are critical for both cortical and cancellous bone maintenance in male mice. In female mice, sex hormones protect tibial cortical bone but play a minor role in preserving cancellous bone mass in vertebrae and tibiae.

Moreover, this sex hormone withdrawal-induced bone loss in young adult mice was mediated through Ot-ERβ. ORX-caused bone loss in tibial cortical bone (Ct. Ar and Imin) in male LC mice did not appear in ORX mice with Ot-ERβ deletion, showing that the lack of Ot-ERβ blunted cortical bone against ORX-induced bone loss in male mice. In contrast, cancellous bone loss in L4 and tibiae by ORX showed no difference between genotypes. It has been shown in previous studies that both testosterone and estrogen protect the cortical bone in male mice [17,18,25,30], suggesting that Ot-ERβ might be sensitive to the deficiency of estrogen and testosterone in young adult male mice to mediate the cortical bone loss, but may not play relevant roles in cancellous bone. Moreover, it was shown that Ot-ERβ was required for cortical bone maintenance in the gonadal intact male mice, which further confirms that Ot-ERβ mediates the protective effect of sex hormones on cortical bone in male mice. Compared to male mice, OVX-induced decreases in tibial cortical and cancellous bone morphology were not influenced by the absence of Ot-ERβ, suggesting that Ot-ERβ may not be involved in the sex hormone regulations of skeletal maintenance in female mice. Thus, we conclude that Ot-ERβ plays a sex-specific role in mediating the sex hormone regulations of skeletal protection. Ot-ERβ is required for cortical bone protection by sex hormones in male mice, whereas it may not mediate sex hormone regulations of cancellous bone in male and female mice.

There are two subtypes of estrogen receptors (ERs), ERα and ERβ, both expressed by all bone cell types in humans and rodents [1,15,53,54]. ERs play important roles in skeletal maintenance in both males and females through binding to estrogen [2,4,55–59]. It is also known that testosterone can activate ERs through direct ligand-binding or aromatizing into estrogen [4,15,25]. A previous study showed that bone loss by ORX was partially protected by testosterone treatment in male mice with global AR knockout (KO) [60], suggesting that ERs might be responsible for this testosterone effect on skeletal protection in male mice. In addition, ERs can also bind to testosterone by forming ER/AR heterodimers [4,28,61]. In earlier studies on young adult and aged male mice with global ERs KO, testosterone was implicated in protecting bone mass through ERβ and AR but independently of ERα [25,30]. Despite using global KO models, these studies still provide evidence for our conjectures that Ot-ERβ regulates cortical bone protection in male mice through binding to testosterone, a process may be associated with ERβ/AR interaction.

In contrast, the ORX-induced cancellous bone loss in male mice was not altered by the lack of Ot-ERβ, suggesting that this cancellous bone protection by sex hormones may be mediated through ERα. ERα in osteoblasts and osteocytes has been demonstrated to mediate the cancellous bone morphology in gonadal intact male mice [31,62]. Estradiol treatments for ORX mice with global ERα KO showed that ERα is required for the estrogen protection of cancellous bone in young and adult male mice (14wk and 28wk) [25,30,59]. In addition, global dual KOs of ERα and ERβ in ORX mice further suggested that ERα and AR, but not ERβ, protect cancellous bone in male mice in response to estrogen [18,25,28] and androgens (testosterone/DHT) [18,25], respectively. Therefore, both ERα and AR, rather than ERβ, are responsible for the sex hormone regulations of cancellous bone in male mice. It is necessary to notice that the elevated circulating sex hormones in global KO models would cause new bone formation before the gonadectomy surgeries [55,63,64], which might magnify skeletal response to the sex hormone deficiency or reduce the effect of the hormonal administrations on bone mass recovery. However, studies on ORX mice with conditional ERs KOs are lacking. Our study is the first to reveal the role of Ot-ERβ in skeletal response to sex hormone deficiency in male mice. Combining with the previous studies, we conclude that in male mice, Ot-ERβ is required for cortical bone maintenance in response to both estrogen and testosterone. In contrast, ERα may primarily mediate the estrogen effect on cortical bone maintenance in male mice. Further work on conditional KOs of ERs and AR in ORX mice with sex hormonal treatments is necessary to elucidate the sex hormone-ERs, particularly the testosterone-ERβ pathway and ERβ/AR interactions in male mice skeletons.

Compared to ERβ, ERα is suggested to be more sensitive to estrogen [25,65–67], which implies that the estrogen effects on skeleton protection in females are primarily modulated through ERα. The osteoprotective role of ERα in osteoblasts and osteocytes has been widely demonstrated in gonadal intact female mice with conditional KO [43,62,68,69]. Moreover, estrogen treatment in OVX mice with global ERα KO showed that the osteoprotective regulation of estrogen in female mice is mainly through ERα [25,29]. Conditional ERα KO in osteocytes and osteoclasts (Oc) of OVX mice further revealed that Ot-ERα and Oc-ERα are primarily responsible for the estrogen protection in trabecular and cortical bone for female mice compared to ERβ [31,70], which explains the similar bone loss we observed between ERβ-dOT and LC mice response to OVX. Consistent with our findings, it is also shown in osteoprogenitor progenitor cells (Ob) that ERβ does not alter the OVX-induced bone loss [71], which further revealed the dispensable role of Ob/Ot-ERβ in mediating the estrogen regulations of female mice skeletons. Collectively, it is indicated that in female mice, Ot-ERα, rather than Ot-ERβ or Ob-ERβ, is crucial for the estrogen response in cortical and cancellous bone.

The morphology and quality of bone are significant contributors to skeletal strength. Bone loss alters the size and geometry of cortical and cancellous bone, resulting in the reduction of mechanical properties and the overall structural strength of the bone, which increases the risk of bone damage and fracture [2,4,32]. Therefore, as an essential factor for bone mass maintenance, sex hormones also play crucial roles in protecting the skeletal strength of males and females [4,49]. Bone loss induced by the sex hormone withdrawal altered the tibial mechanical environment, as reflected by increased FE-predicted principal peak strains. These strain changes are consistent with reduced mechanical competence. Specifically, ORX-induced bone loss in the tibia led to the reduced mechanical strength of both tibial cortical and cancellous bone, resulting in a decreased stiffness of the overall tibia in male LC mice compared to the Sham mice. In female LC mice with OVX, greater tensile and compressive strains appeared in tibial cancellous but not cortical bone, whereas the cancellous bone morphology at the proximal tibia was not altered by the sex hormone deficiency. This increased tibial cancellous bone strain in the OVX LC mice may be associated with the loss of the cortical shell around the proximal trabecular bone region. The thinning of the cortical shell may compromise its load-bearing ability, which causes the inner trabecular bone to carry more loads. OVX-induced decreases in cortical bone mineral density and Imax at the proximal tibia (37%) indicate similar bone loss in the cortical bone shell at the metaphyseal tibia. Further morphological analysis is necessary to offer more specific evidence to support this conjecture.

Moreover, the sex-specific role of Ot-ERβ in skeletal morphology by sex hormones is also mirrored in the tibial strength in male and female mice. Consistent with the morphological results, deletion of Ot-ERβ prevented ORX-induced increases in the cortical and cancellous bone strains and protected the stiffness of the overall tibia, suggesting that Ot-ERβ mediates the bone loss-caused reduction in tibial strength in male mice in response to the sex hormone deficiency. Meanwhile, it needs to be noted that in the gonadal intact male mice lacking Ot-ERβ, cancellous bone, with unaltered bone mass, showed higher tensile and compressive strains than LC mice. These increases in mechanical strains could be caused by structural and geometric changes [72,73]. The orientation and architecture of trabeculae may vary with the deletion of Ot-ERβ and, therefore, reduce mechanical strength. More analysis on the trabecular configuration needs to be done to provide a reasonable explanation.

In contrast, the deletion of Ot-ERβ in female mice did not alter the effect of OVX on peak strains, neither tensile nor compressive, in tibial cortical and cancellous bone. The lack of sex hormones had no impact on the peak strains in the tibial cortical bone of female LC and ERβ-dOT mice, matching the unaltered cortical bone morphology. However, despite a similar bone mass, tibial cortical bone in the gonadal intact female mice with Ot-ERβ deletion experienced lower peak strains, associated with greater whole-bone stiffness than the LC mice. The higher tibial strength in the female mice lacking Ot-ERβ may be related to the longitudinal curvature or the shape of the cortical bone, as the deletion of Ot-ERβ in gonadal intact female mice results in a more vertical shape of the tibia relative to the LC mice, allowing the proximal and midshaft cortical bone to bear higher loads. A more detailed analysis of the curvature of the tibial cortical bone is needed to validate this conjecture.

Also, future studies with administrations of estradiol and testosterone for the ORX/OVX mice will help explain the interaction of sex hormones and Ot-ERβ in male and female mice on bone mass maintenance. An additional limitation in this study would be the lack of histological analysis, which may offer more evidence on bone cell activities to explain the sex hormone regulations of bone turnover.

Collectively, this study revealed the critical roles of sex hormones in protecting bone mass and skeletal strength in both young adult male and female mice. Moreover, we revealed the role of Ot-ERβ in mediating the effects of sex hormone deficiency on skeletal strength in male and female mice, which matches its relevant regulations of bone morphology. The protective effects of sex hormones on cortical but not cancellous bone morphology in male mice are mediated by Ot-ERβ. That lack of Ot-ERβ in male mice blunts sex hormone deficiency-induced reductions of cortical bone morphology and load-induced peak strains in cortical and cancellous bone and the overall strength of the tibia. In contrast, sex hormones are essential for maintaining tibial cortical bone but may not mediate cancellous bone mass in female LC mice. Moreover, Ot-ERβ is not required for the OVX-induced changes in the bone mass or skeletal strength of cortical and cancellous bone in female mice.

## Acknowledgements

This work was completed under the mentorship of the *late* Dr. Russell P. Main, whose scientific guidance, intellectual insight, and commitment to rigorous biomechanics research were essential to the conception, execution, and interpretation of this study. The authors are deeply grateful for his mentorship and lasting contributions to this work.

We thank members of the Musculoskeletal Biology and Mechanics Laboratory at Purdue University for helpful discussions and technical support. We also acknowledge the animal care staff and research support personnel at Purdue University for their assistance with mouse colony maintenance, surgical procedures, imaging, and sample processing. This work was supported by NIH grant F32 AR054676.

---

**Supplementary Table S1.**
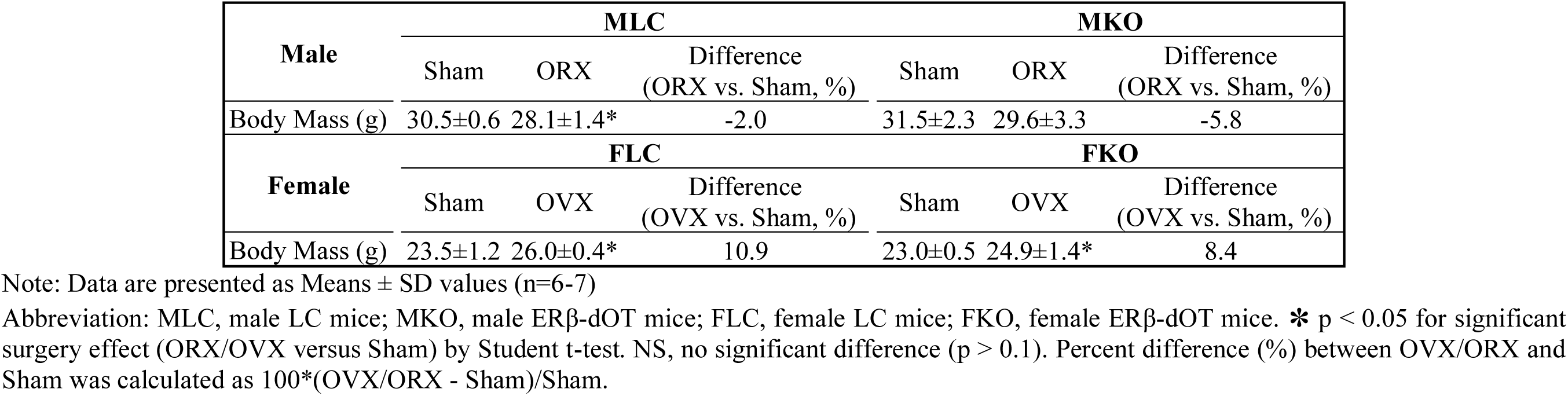
Body mass of the male and female mice ERβ-dOT and LC mice with gonadectomy (ORX/OVX) or Sham at 24 wks of age.

**Supplementary Table S2.**
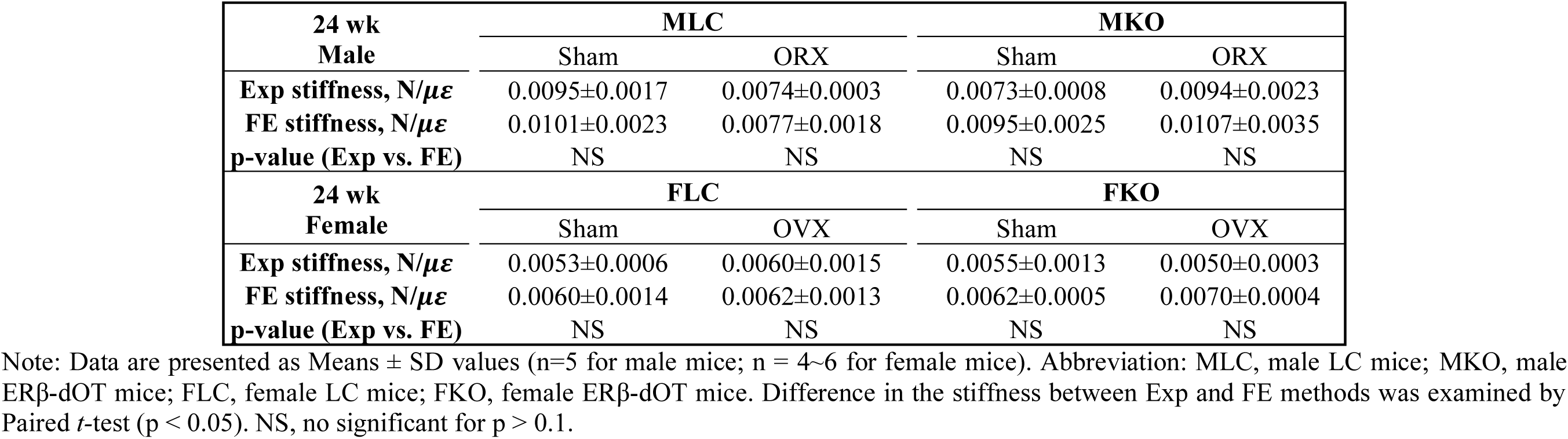
Stiffness measurements of the gauge-based experiment (Exp) and finite element modeling (FE) at the gauge site for male and female mice ERβ-dOT and LC mice with gonadectomy (ORX/OVX) or Sham at 24 wks of age.

| 24 wk<br>Male | MLC |  | MKO |  |
| --- | --- | --- | --- | --- |
|  | Sham | ORX | Sham | ORX |
| Exp stiffness, N/$\mu$ $\epsilon$ | 0.0095±0.0017 | 0.0074±0.0003 | 0.0073±0.0008 | 0.0094±0.0023 |
| FE stiffness, N/$\mu$ $\epsilon$ | 0.0101±0.0023 | 0.0077±0.0018 | 0.0095±0.0025 | 0.0107±0.0035 |
| p-value (Exp vs. FE) | NS | NS | NS | NS |
| 24 wk<br>Female | FLC |  | FKO |  |
|  | Sham | OVX | Sham | OVX |
| Exp stiffness, N/$\mu$ $\epsilon$ | 0.0053±0.0006 | 0.0060±0.0015 | 0.0055±0.0013 | 0.0050±0.0003 |
| FE stiffness, N/$\mu$ $\epsilon$ | 0.0060±0.0014 | 0.0062±0.0013 | 0.0062±0.0005 | 0.0070±0.0004 |
| p-value (Exp vs. FE) | NS | NS | NS | NS |

Note: Data are presented as Means ± SD values (n=5 for male mice; n = 4~6 for female mice). Abbreviation: MLC, male LC mice; MKO, male ERβ-dOT mice; FLC, female LC mice; FKO, female ERβ-dOT mice. Difference in the stiffness between Exp and FE methods was examined by Paired *t*-test ( $p < 0.05$ ). NS, no significant for $p > 0.1$ .

**Supplementary Table S3.**
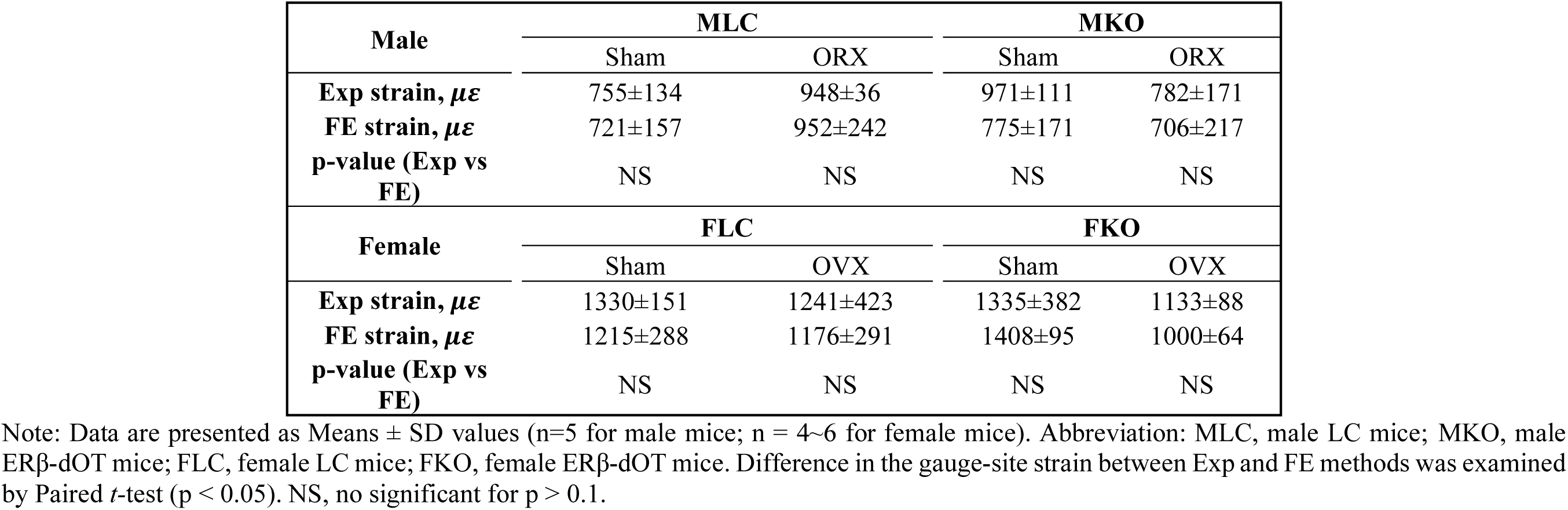
Gauge-measured (Exp) and FE-predicted (FE) strains (*μɛ*) at the gauge site under −7N applied load in male and female mice ERβ-dOT and LC mice with gonadectomy (ORX/OVX) or Sham at 24 wks of age

| Male | MLC |  | MKO |  |
| --- | --- | --- | --- | --- |
|  | Sham | ORX | Sham | ORX |
| Exp strain, $\mu\epsilon$ | 755±134 | 948±36 | 971±111 | 782±171 |
| FE strain, $\mu\epsilon$ | 721±157 | 952±242 | 775±171 | 706±217 |
| p-value (Exp vs FE) | NS | NS | NS | NS |
| Female | FLC |  | FKO |  |
|  | Sham | OVX | Sham | OVX |
| Exp strain, $\mu\epsilon$ | 1330±151 | 1241±423 | 1335±382 | 1133±88 |
| FE strain, $\mu\epsilon$ | 1215±288 | 1176±291 | 1408±95 | 1000±64 |
| p-value (Exp vs FE) | NS | NS | NS | NS |

Note: Data are presented as Means ± SD values (n=5 for male mice; n = 4~6 for female mice). Abbreviation: MLC, male LC mice; MKO, male ERβ-dOT mice; FLC, female LC mice; FKO, female ERβ-dOT mice. Difference in the gauge-site strain between Exp and FE methods was examined by Paired *t*-test ( $p < 0.05$ ). NS, no significant for $p > 0.1$ .

