## Supplementary Tables for "The Role of Osteocyte Estrogen Receptor β in Bone Mass Maintenance and Tibial Stiffness Following Sex Hormone Withdrawal in Male and Female Mice": OX-FEA-Supple Figures_062026.docx

**Supplementary Table S1** Body mass of the male and female mice ERβ-dOT and LC mice with gonadectomy (ORX/OVX) or Sham at 24 wks of age

| **Male** |  | **MLC** | | |  | **MKO** | | |
| --- | --- | --- | --- | --- | --- | --- | --- | --- |
|  |  | Sham | ORX | Difference   (ORX vs. Sham, %) |  | Sham | ORX | Difference   (ORX vs. Sham, %) |
| Body Mass (g) |  | 30.5±0.6 | 28.1±1.4* | -2.0 |  | 31.5±2.3 | 29.6±3.3 | -5.8 |
| **Female** |  | **FLC** | | |  | **FKO** | | |
|  |  | Sham | OVX | Difference   (OVX vs. Sham, %) |  | Sham | OVX | Difference   (OVX vs. Sham, %) |
| Body Mass (g) |  | 23.5±1.2 | 26.0±0.4* | 10.9 |  | 23.0±0.5 | 24.9±1.4* | 8.4 |

Note: Data are presented as Means ± SD values (n=6-7)

Abbreviation: MLC, male LC mice; MKO, male ERβ-dOT mice; FLC, female LC mice; FKO, female ERβ-dOT mice. ✽ p < 0.05 for significant surgery effect (ORX/OVX versus Sham) by Student t-test. NS, no significant difference (p > 0.1). Percent difference (%) between OVX/ORX and Sham was calculated as 100*(OVX/ORX - Sham)/Sham.

**Supplementary Table S2** Stiffness measurements of the gauge-based experiment (Exp) and finite element modeling (FE) at the gauge site for male and female mice ERβ-dOT and LC mice with gonadectomy (ORX/OVX) or Sham at 24 wks of age

| **24 wk Male** |  | **MLC** | |  | **MKO** | |
| --- | --- | --- | --- | --- | --- | --- |
|  |  | Sham | ORX |  | Sham | ORX |
| **Exp stiffness, N/**$\boldsymbol{\mu\varepsilon}$ |  | 0.0095±0.0017 | 0.0074±0.0003 |  | 0.0073±0.0008 | 0.0094±0.0023 |
| **FE stiffness, N/**$\boldsymbol{\mu\varepsilon}$ |  | 0.0101±0.0023 | 0.0077±0.0018 |  | 0.0095±0.0025 | 0.0107±0.0035 |
| **p-value (Exp vs. FE)** |  | NS | NS |  | NS | NS |
| **24 wk Female** |  | **FLC** | |  | **FKO** | |
|  |  | Sham | OVX |  | Sham | OVX |
| **Exp stiffness, N/**$\boldsymbol{\mu\varepsilon}$ |  | 0.0053±0.0006 | 0.0060±0.0015 |  | 0.0055±0.0013 | 0.0050±0.0003 |
| **FE stiffness, N/**$\boldsymbol{\mu\varepsilon}$ |  | 0.0060±0.0014 | 0.0062±0.0013 |  | 0.0062±0.0005 | 0.0070±0.0004 |
| **p-value (Exp vs. FE)** |  | NS | NS |  | NS | NS |

Note: Data are presented as Means ± SD values (n=5 for male mice; n = 4~6 for female mice). Abbreviation: MLC, male LC mice; MKO, male ERβ-dOT mice; FLC, female LC mice; FKO, female ERβ-dOT mice. Difference in the stiffness between Exp and FE methods was examined by Paired *t-*test (p < 0.05). NS, no significant for p > 0.1.

**Supplementary Table S3** Gauge-measured (Exp) and FE-predicted (FE) strains ($\mu\varepsilon$ ) at the gauge site under -7N applied load in male and female mice ERβ-dOT and LC mice with gonadectomy (ORX/OVX) or Sham at 24 wks of age

| **Male** |  | **MLC** | |  | **MKO** | |
| --- | --- | --- | --- | --- | --- | --- |
|  |  | Sham | ORX |  | Sham | ORX |
| **Exp strain,** $\boldsymbol{\mu\varepsilon}$ |  | 755±134 | 948±36 |  | 971±111 | 782±171 |
| **FE strain,** $\boldsymbol{\mu\varepsilon}$ |  | 721±157 | 952±242 |  | 775±171 | 706±217 |
| **p-value (Exp vs FE)** |  | NS | NS |  | NS | NS |
| **Female** |  | **FLC** | |  | **FKO** | |
|  |  | Sham | OVX |  | Sham | OVX |
| **Exp strain,** $\boldsymbol{\mu\varepsilon}$ |  | 1330±151 | 1241±423 |  | 1335±382 | 1133±88 |
| **FE strain,** $\boldsymbol{\mu\varepsilon}$ |  | 1215±288 | 1176±291 |  | 1408±95 | 1000±64 |
| **p-value (Exp vs FE)** |  | NS | NS |  | NS | NS |

Note: Data are presented as Means ± SD values (n=5 for male mice; n = 4~6 for female mice). Abbreviation: MLC, male LC mice; MKO, male ERβ-dOT mice; FLC, female LC mice; FKO, female ERβ-dOT mice. Difference in the gauge-site strain between Exp and FE methods was examined by Paired *t-*test (p < 0.05). NS, no significant for p > 0.1.
